# Generating New Musical Preferences from Multi-level Mapping of Predictions to Reward

**DOI:** 10.1101/2022.06.17.496615

**Authors:** Nicholas Kathios, Matthew E. Sachs, Euan Zhang, Yongtian Ou, Psyche Loui

## Abstract

Much of what we know and love about music hinges on our ability to make successful predictions, which appears to be an intrinsically rewarding process. Yet the exact process by which learned predictions become pleasurable is unclear. Here, we created novel melodies in an alternative scale different from any established musical culture, to show how musical preference is generated *de novo*. Across nine studies (n=1185), participants learned to like more frequently-presented items that adhered to this rapidly-learned structure, suggesting that exposure and prediction errors both affected self-report liking ratings. Learning trajectories varied by music reward sensitivity, but were similar for USA and Chinese participants. Furthermore, fMRI activity in auditory areas reflected prediction errors whereas functional connectivity between auditory and medial prefrontal regions reflected both exposure and prediction errors. Collectively, results support predictive coding as a cognitive mechanism by which new musical sounds become rewarding.

## Statement of Relevance

Here we address the question of why humans find music listening to be psychologically rewarding by combining large-scale behavioral testing, cross-cultural experiments, neuropsychological testing, functional MRI, and music theory and music technology. Borrowing from theories in statistical learning, predictive coding, and reward learning, in nine studies we systematically manipulate exposure and prediction error, to establish prediction to reward learning trajectories that are sensitive to individual differences but relatively stable across cultures, and that are tied to the activity and functional connectivity of the auditory and reward systems of the brain.

## Introduction

Why do we love music? In contrast to other pleasures in life, such as food and sex, music has no obvious adaptive value; yet an attraction to music is ubiquitous across cultures and across the lifespan. Indeed, both listening to and performing music ranks highly among life’s greatest pleasures [1], and reliably engages the dopaminergic reward system [2–4]. Classic work [48, 49] has posited a relationship between arousal and aesthetic preference, resulting in effects of novelty, familiarity, and mere exposure (ME) on preference [50, 18]. Recent findings have supported an adjacent view, that of predictive coding (PC): as predictions and reward signals are ubiquitous features of the central nervous system that underlie perception, action, and emotion [5–8], so too may the rewarding effects of music listening come from making successful predictions and minimizing prediction errors [9, 10].

These accounts, while ostensibly similar, make subtly different predictions. A classic ME-based account posits that repeated exposure to sound sequences with predictable statistical probabilities can change preferences for those specific sound sequences [17], resulting in a classic inverted-u model of preference to music as a function of familiarity and novelty [18]. A PC-based account posits that preference comes from making successful predictions and minimizing prediction errors [10]. This model is motivated in part by work on musical expectations, which proposes that musical expectations can form from learning statistical regularities and patterns in music (schematic expectations) as well as familiarity with a particular piece of music or genre of music (veridical expectations); [48, 10, 11]). Thus, while the ME-based model would predict that preferences for novel sequences are informed only by the fact that the sequences have not been heard before (thus generating little veridical expectations for these sequences), the PC model would predict that preferences would be informed by the degree to which listeners can minimize the prediction errors that these sequences elicit. For instance, shared features across sound sequences, such as common harmonic or melodic structures that aid in minimizing prediction errors, would lead to greater preferences for these sequences, according to the PC model.

Beyond sequence-specific (veridical) knowledge, predictions can unfold at multiple levels, affecting schematic expectations for music, whether they be stylistic (hip-hop, jazz), structural (melody, tonality), temporal (rhythm, meter), and/or acoustic (pitch, timbre) [11, 12, 51, 52]. This offers several ways by which novel music, for which there is no veridical knowledge, can nonetheless confirm listeners’ predictions. Evaluating ME and PC accounts against each other requires the ability to manipulate both exposure and prediction error: exposure by presenting musical items a variable number of times, and prediction errors by presenting musical items that adhere to (or not adhere to) a particular musical structure at varying levels of exposure Furthermore, as the PC account draws from models of dopaminergic function [5–8], measuring activity in the reward system of the brain with fMRI would provide a strong test of our ability to manipulate exposure and prediction.

While it has been shown the precise relationships between exposure, prediction error, and reward may vary across cultures [53] and/or with individual differences in reward sensitivity to music [54], it is often challenging to understand how exposure relates to learning and reward due to the fact that most stimuli that we encounter, even for the first time, makes use of overlearned predictions to which we may have been exposed throughout our lives. This is especially the case with musical structures, such as common sets of pitches or musical scales that we have implicitly acquired from lifelong exposure [20]. As a concrete example of such knowledge, most listeners within Western cultures show implicit knowledge of, and preference for, common-practice Western musical scale structures based around the octave, which is a doubling of acoustic frequency [21]. We circumvent this challenge of overlearned predictions by incorporating a unique and unfamiliar musical system: the Bohlen-Pierce (B-P) scale, which is based on a tripling of acoustic frequency, thus differing acoustically and statistically from the world’s existing musical systems [22].

Here we test the effects of exposure and prediction error on musical learning and preference by using naturalistic music composed in grammatical structures defined in the B-P scale [17]. In Studies 1-4, we ask the degree to which self-reported familiarity and liking ratings reflect exposure and prediction error. Exposure was manipulated by presenting novel naturalistic musical melodies a variable number of times, and prediction error was manipulated through structural alterations to the endings of the exposed melodies. In Study 5, we test these relationships in cases of congenital and acquired music anhedonia. In Study 6, we reversed the presentation of altered and original sets of melodies to tease apart differential contributions of schematic and veridical expectations to familiarity and liking ratings. In Study 7, we ensure that results from previous studies are not due to anchoring effects. In Study 8, we test the effects of culture on predictions and reward in a cross-cultural replication on a sample from China. Finally, in Study 9, we evaluate effects of this learning on reward system activity and connectivity using fMRI. Together, the studies trace the trajectory of preference learning from exposure to melodic and statistical structures in a novel musical system. As the human ability to recognize and learn statistical properties of stimuli via mere exposure has been posited to underlie multiple cognitive tasks beyond music, including language acquisition [13, 14] and decision-making [15], our results provide a mechanistic account not only for why people enjoy music, but also the circumstances under which our ability to predict leads to reward, a concept that underlies much of motivated behavior [19].

## Results

### Analysis Plan

For all studies, participants provided familiarity and liking ratings for melodies composed in a predefined grammatical structure (based on the B-P scale [17]) that were either 1) presented a variable number of times in an exposure phase (*effect of exposure*), or 2) altered to have a different, non-grammatical ending than the original melodies that were presented during exposure (*effect of prediction error*). In each study, only one set of melodies was presented during the exposure phase (“original” melodies), while the other set of melodies contained previously unexposed endings and therefore generated a prediction error (“altered” melodies). Both groups of melodies were rated on both familiarity and liking in a post-exposure rating phase. To investigate the effects of exposure and prediction error on these post-exposure familiarity and liking ratings, we constructed linear mixed-effect models using the R package *lme4* [23]. We included prediction error (original [no prediction error elicited] vs. altered [prediction error elicited]) as an interaction term in these models. This term was effect-coded such that the main effect of exposure represents the average effect across both types of melodies. This modeling allowed us to investigate 1) the main effect of exposure, 2) the main effect of prediction error, and 3) the difference in the effect of prediction error as a function of exposure (the interaction term in the model). Crucially, this interaction term also allowed us to test competing predictions between the ME and PC accounts. Because the melodies presented most often in the exposure phase would be the most predictable, the altered counterparts of these melodies will elicit greater prediction errors than those complementary to the melodies presented less often in the exposure phase. As a result, if preference is informed by the degree to which participants can minimize prediction errors, then the effect of prediction error should be greater for melodies presented more often in the exposure phase (i.e. there will be an interaction).

Conversely, if preference is fundamentally informed by mere exposure, then the effect of prediction errors should remain consistent across the number of presentations (i.e. there will be no interaction). We specified by-participant random slopes (including the interaction term) and intercepts and by-item (melody) random intercepts. Continuous predictor and dependent variables were standardized before being entered into the model. Significance of fixed effects (exposures and prediction error) was determined using the Satterthwaite method to approximate the degrees of freedom with the *lmerTest* package [24].

### Study 1

Participants listened to 8 monophonic musical melodies composed in the B-P scale during the exposure phase. The number of presentations varied for each melody (either 2, 4, 8, or 16 times with two melodies in each condition). After exposure, participants made familiarity and liking ratings for each melody, along with two melodies not heard in the exposure phase (thus, presented 0 times during exposure), as well as altered versions of the 10 melodies, which were identical except for an unexpected ending. For familiarity ratings, there was a significant interaction between exposure and prediction error n (β = 0.01, t(1883) = 3.99, p < 0.001): the effect of prediction error (main effect: β = 0.16, t(1169) = 6.33, p < 0.001) increased as a function of exposure (main effect: β = 0.33, *t*(171) = 21.23, *p* < 0.001). For liking ratings, there was also a significant interaction between exposure and prediction error (β = 0.05, *t*(1200) = 2.27, *p* = 0.02): the effect of prediction error on liking ratings (main effect: β = 0.11, *t*(1793) = 4.94, *p* < 0.001) also increased as a function of exposure (main effect: β = 0.03, *t*(1169) = 2.05, *p* = 0.04). Thus, both exposure and prediction errors informed both familiarity and liking ratings, as participants reported more preference and familiarity for melodies that were both exposed more often and that did not elicit a prediction error. However, these results are more consistent with predictions of the PC model, such that the effect of prediction error on liking ratings increased with the magnitude of the error.

### Study 2

In Study 2, we extended the findings from Study 1 to determine the degree to which changing the specific numbers of presentations during the exposure phase affected liking ratings. In a new group of participants, we replicated Study 1 but with melodies that were presented either 0, 2, 4, 6, 10, or 14 times. For familiarity ratings, there was again a significant interaction between exposure and prediction error (β = 0.06, *t*(3411) = 2.81, *p* = 0.005): again, the effect of prediction error (main effect β = 0.14, *t*(1545) = 5.87, *p* < 0.001) increased as a function of exposure (main effect β = 0.3, t(163) = 15.71, p < 0.001). For liking ratings,we again found a significant main effect of number of presentations (β = 0.02, *t*(163) = 2.1, p = 0.04) and prediction error (β = 0.02, *t*(171) = 2.1, *p* < 0.001). However, we did not detect an interaction between prediction error and exposure(β = 0.13, *t*(3179) = 1.06, *p* = 0.29). As the sample size of these studies were chosen to detect the effect of exposure rather than to detect an interaction (see Materials and Methods), the lack of interaction could simply be due to insufficient statistical power; thus we went on to replicate and extend these studies and to test for an interaction with aggregated data across several studies.

### Studies 3 and 4

Studies 3 and 4 were designed to replicate the findings from Studies 1 and 2 with a new sample. Study 3 used the same numbers of presentation as Study 1 (0, 2, 4, 8, 16) and Study 4 used the same numbers of presentation as Study 2 (0, 2, 4, 6, 10, 14). For familiarity ratings in Study 3, there was a significant interaction between exposure and prediction error (β = 0.07, *t*(2305) = 2.79, *p* = 0.005): again, the effect of prediction error (β = 0.14, *t*(1168) = 5.81, *p* < 0.001) increased as a function of number of presentations (main effect β = 0.33, *t*(168) = 18.81, *p* < 0.001). For liking ratings, we also replicated the main effect of exposure (β = 0.06, t(169) = 3.66, p < 0.001). Again, melodies that did not elicit a prediction error were preferred over melodies with prediction errors (β = 0.07, *t*(1507) = 3.15, *p* = 0.002). There was no interaction between the two (β = 0.01, *t*(1434) = 0.56, *p* = 0.57). For familiarity ratings in Study 4, we replicated the main effect of exposure (β = 0.34, *t*(163) = 19.62, *p* < 0.001) and prediction error (β = 0.17, *t*(1923) = 7.08, *p* < 0.001). We did not detect a prediction error X exposure interaction (β = 0.04, *t*(2026) = 1.66, *p* = 0.1). For liking ratings in Study 4, we replicated the significant effect of exposure (β = 0.03, *t*(162) = 2.14, *p* = 0.03). Melodies that did not elicit a prediction error were once again rated as more liked than melodies that did (β = 0.09, *t*(3316) = 4.67, *p* < 0.001). There was no interaction between prediction error and exposure (β = 0.02, *t*(1801) = 0.87, *p* = 0.38). Together, these four studies consistently show that main effects of exposure and prediction error were robust for both familiarity and liking, but the interaction was much more variable, especially for liking. Since Studies 1 through 4 used different samples of participants but the same stimuli with different numbers of presentations, we proceeded to combine the data from these studies for a mini meta-analysis to evaluate the effects of, and interaction between, prediction error and exposure on familiarity and liking across a larger sample.

### Mini Meta-Analyses of Studies 1-4

#### Familiarity ratings shows a logarithmic relationship with exposure

When considering the shape of the relationship between exposure and familiarity, we expected that familiarity ratings would show a logarithmic relationship with exposure (i.e. participants would learn the stimuli after a certain amount of presentations, after which subsequent presentations do not make them more familiar), as opposed to a more linear relationship (i.e. ratings continue to increase with the exposure). We compared the fit between logarithmic and linear models for combined data across Studies 1-4 (n = 667). These models had the same random effects structure as previous models. Results from this mini meta-analysis showed both main effects of number of presentations and alterations, as well as significant interactions between alterations and number of presentations, in both linear and logarithmic models. Following the suggestion of Zuur et al. [25], parameters were estimated using maximum likelihood to enable model comparison, and Akaike’s Information Criteria (AIC) was compared across these models to compare their fit. This revealed that a logarithmic model (AIC = 31575) was a better fit compared to a linear model (AIC = 33986) to model the relationship between number of presentations and familiarity ratings (see Table 1 and Figure 1).

**Table 1.**
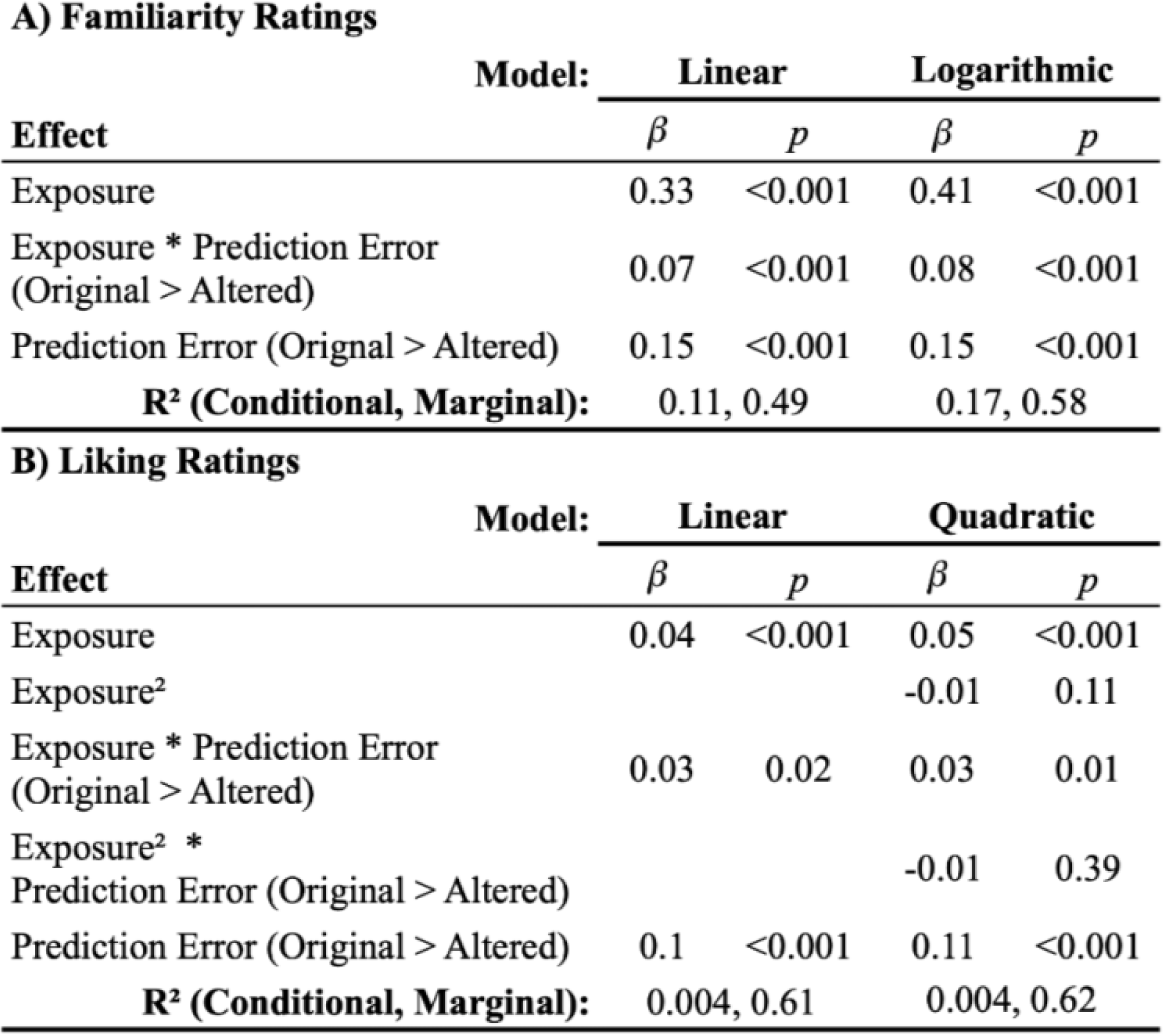
Standardized Beta coefficients, associated p-values, and R^2^ values for each model fit for familiarity (A) and liking (B) ratings.

**Figure 1.**
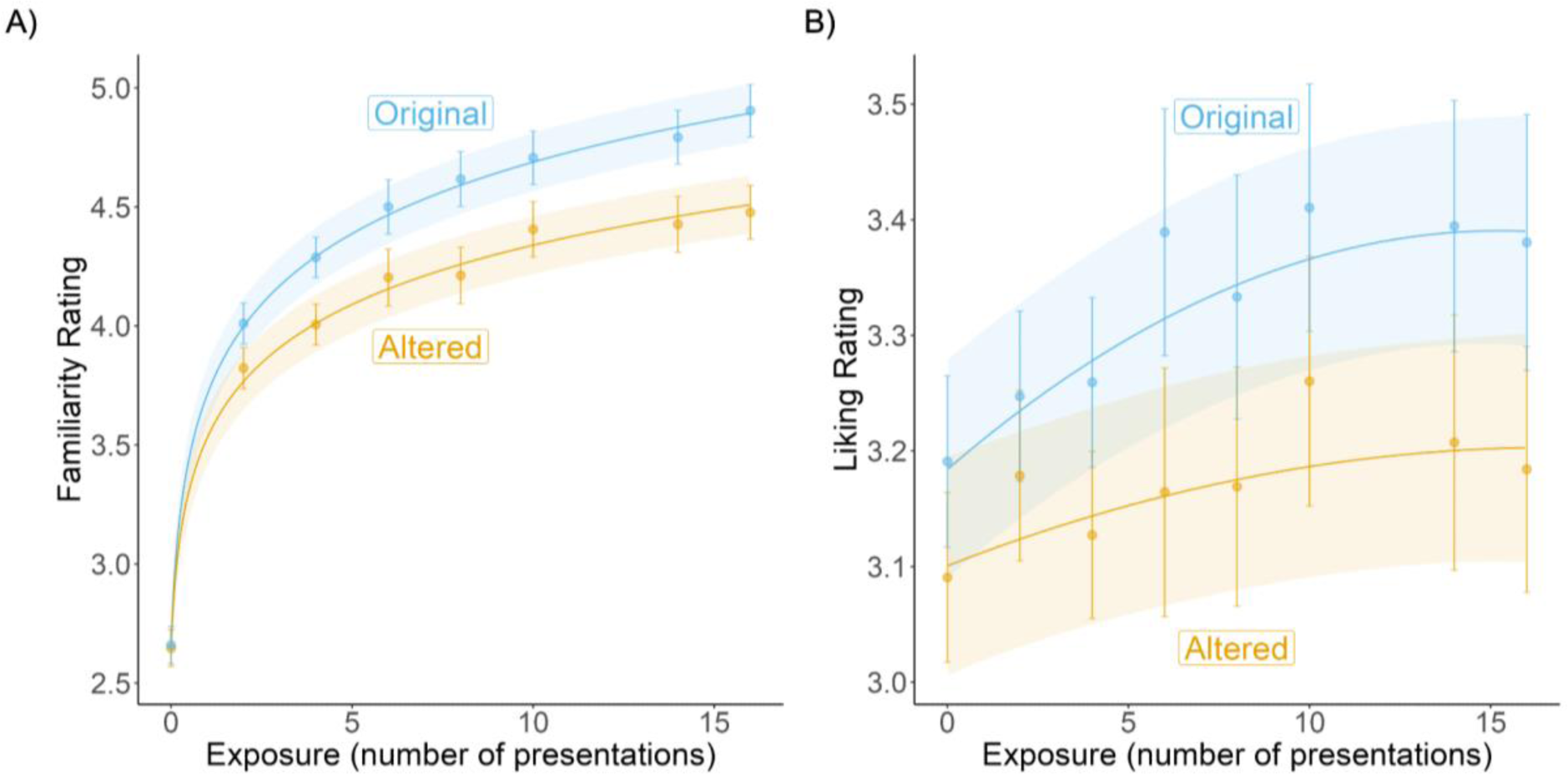
Best-fit model predictions of familiarity (A) and liking (B) ratings as a function of exposure and prediction error (“Original” melodies did not elicit a prediction error while “Altered” melodies did) across Studies 1 to 4. Points and associated error bars indicate mean ratings and 95% confidence intervals.

#### Liking ratings show a quadratic relationship with exposure

We used the same approach to best describe the relationship between liking ratings and exposure. However, as the trajectory between exposure and liking typically shows an inverse-U relationship (see Chmiel & Schubert [18] for a review), we compared model fits of a linear and quadratic model using a Likelihood Ratio test. Both linear and quadratic models showed significant main effects of number of presentations and alterations, as well as significant interactions between the two. The quadratic model was found to best describe the relationship between number of presentations and liking ratings (χ^2^(13) = 127.03, *p* < 0.001; see Table 1 for model fits and Figure 1 for model predictions plotted with mean and 95% confidence intervals). Together, these results are most consistent with the predictions of the PC model: the fact that the effect of prediction error increased as a function of exposure suggests that the minimization of prediction errors best accounts for the generation of musical preferences in the context of this novel musical environment.

#### Music reward sensitivity influences the learning trajectory

While results from Studies 1-4 showed musical preferences were informed by both exposure and prediction error across an aggregated sample of 667 participants, past work has also shown considerable individual differences in music reward sensitivity, which affects the degree to which individuals enjoy music listening [26]. Thus, we tested the hypothesis that liking ratings of individuals with low music reward sensitivity, i.e. musical anhedonics, would show decreased sensitivity to our manipulations of both exposure and prediction error. Following past work [26], we split our aggregated sample into tertiles using the Barcelona Music Reward Questionnaire (BMRQ), a measure of music reward sensitivity [27]. These tertiles represent relatively high (hyperhedonics, BMRQ = 86-100), medium (hedonics, BMRQ = 76-85), and low sensitivity (anhedonics, BMRQ = 26-75) to music reward in our sample. To test our hypothesis, we added an interaction term for music reward sensitivity to our best fitting models (logarithmic for familiarity ratings; quadratic for liking ratings). As this measure indexes individual differences in music reward sensitivity, we expected differences across these tertiles only on liking ratings. If musical anhedonics’ liking ratings are less sensitive to exposure effects than their more hedonic counterparts, then they will show a different trajectory between liking ratings and exposure (i.e. a music reward sensitivity X exposure interaction). If they are less sensitive to prediction errors, then they will show a decreased effect of prediction error (i.e. a music reward sensitivity X prediction error interaction). For these analyses, this variable was dummy-coded to treat the hedonic group as the reference level. We interpreted any interaction between number of presentations and music reward sensitivity as evidence that the relationship between familiarity and/or liking ratings and number of presentations differed across groups.

For familiarity ratings, there were no differences in ratings across the three tertiles. Further, there were no significant two-way interactions between music reward sensitivity and exposure or prediction error, and no significant three-way interaction between music reward sensitivity, exposure, and prediction error (see Table 2 for model fits and Figure 2 for model predictions plotted with mean and 95% confidence intervals). This suggests that musical anhedonics familiarize themselves similarly to music compared to their more hedonic counterparts, in that their ratings were similarly sensitive to both exposure and prediction error.

**Table 2.** Standardized Beta coefficients and associated p-values for the three-way (music reward sensitivity X exposure X prediction error) interaction models built on (A) familiarity and (B) liking ratings. Only music reward sensitivity terms are shown.

| A) Familiarity Ratings |  |  |  |
| --- | --- | --- | --- |
| Effect | Tertile Contrast | $\beta$ | $p$ |
| Music Reward Sensitivity | Hyperhedonic > Hedonic | 0.06 | 0.26 |
|  | Hedonic > Anhedonic | 0.02 | 0.66 |
| Music Reward Sensitivity * Exposure | Hyperhedonic > Hedonic | 0.01 | 0.6 |
|  | Hedonic > Anhedonic | -0.008 | 0.75 |
| Music Reward Sensitivity * Prediction Error (Original > Altered) | Hyperhedonic > Hedonic | 0.008 | 0.78 |
|  | Hedonic > Anhedonic | -0.005 | 0.86 |
| Music Reward Sensitivity * Exposure * Prediction Error (Original > Altered) | Hyperhedonic > Hedonic | 0.03 | 0.34 |
|  | Hedonic > Anhedonic | -0.005 | 0.85 |
| R<sup>2</sup> (Conditional, Marginal): |  | 0.17, 0.59 |  |
| B) Liking Ratings |  |  |  |
| Effect | Tertile Contrast | $\beta$ | $p$ |
| Music Reward Sensitivity | Hyperhedonic > Hedonic | 0.06 | 0.4 |
|  | Hedonic > Anhedonic | 0.17 | 0.02 |
| Music Reward Sensitivity * Exposure | Hyperhedonic > Hedonic | 0.02 | 0.47 |
|  | Hedonic > Anhedonic | -0.02 | 0.38 |
| Music Reward Sensitivity * Exposure<sup>2</sup> | Hyperhedonic > Hedonic | -0.03 | 0.14 |
|  | Hedonic > Anhedonic | 0.06 | 0.002 |
| Music Reward Sensitivity * Prediction Error (Original > Altered) | Hyperhedonic > Hedonic | -0.01 | 0.79 |
|  | Hedonic > Anhedonic | 0.03 | 0.37 |
| Music Reward Sensitivity * Exposure * Prediction Error (Original > Altered) | Hyperhedonic > Hedonic | 0.0006 | 0.99 |
|  | Hedonic > Anhedonic | 0.02 | 0.5 |
| Music Reward Sensitivity * Exposure<sup>2</sup> * Prediction Error (Original > Altered) | Hyperhedonic > Hedonic | 0.007 | 0.81 |
|  | Hedonic > Anhedonic | -0.003 | 0.91 |
| R<sup>2</sup> (Conditional, Marginal): |  | 0.02, 0.62 |  |

**Figure 2.**
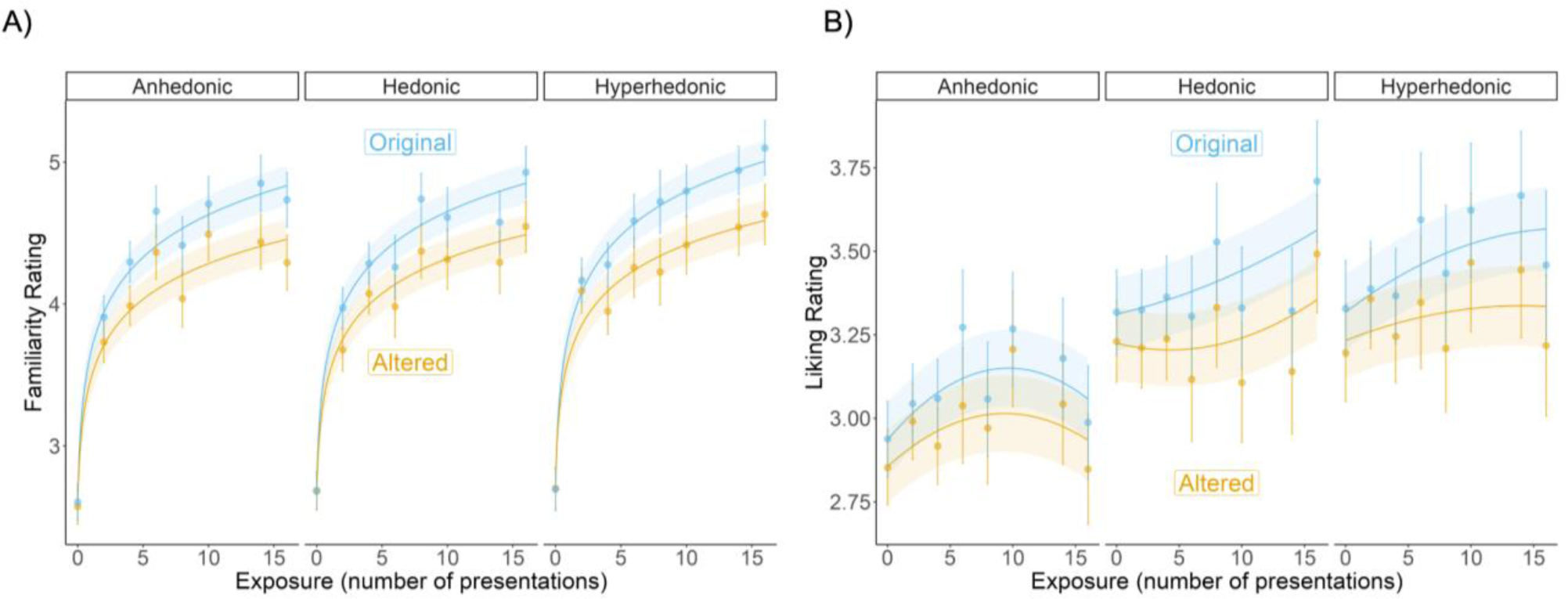
Model-predicted familiarity (A) and liking (B) ratings as a function of exposure, prediction error, and music reward sensitivity (tertile split on BMRQ: anhedonic, hedonic, and hyperhedonic groups). Points and associated error bars represent 95% confidence intervals.

For liking ratings, there was a significant difference across groups: the hedonic group rated melodies as more liked than the anhedonic group (β = 0.17, *t*(663) = 2.36 *p* = 0.02). There were no significant linear interactions between exposure and music reward sensitivity, and there were not any interactions between prediction error and music reward sensitivity. There was no significant three-way interaction between music reward sensitivity, exposure, and prediction error. We did, however, detect an interaction between the quadratic exposure term and music reward sensitivity (interaction β = 0.06, *t*(649) = 3.05, *p* = 0.002): while the anhedonic group showed a significant inverse-U relationship between exposure and liking ratings (β = -0.04, *t*(650) = -3.12 *p* = 0.002), the hedonic and hyperhedonic groups did not (hedonic: β = 0.02, *t*(649) = 0.24; hyperhedonic: β = -0.01, *t*(675) = 0.95, *p* = 0.34) (for all results, see Table 2, Figure 2). These results suggest that, while all groups responded similarly to prediction errors, continued exposure to these melodies led to increased liking in all but the anhedonic group.

### Study 5

While results from the tertile split above show that large online samples can capture a range of musical reward sensitivity that predicts differences in learning to like new music, extreme cases of insensitivity to reward can be effective tests of the models derived from the studies above. In Study 5 we tested the models relating exposure and prediction error to familiarity and liking for the anhedonic, hedonic, and hyperhedonic subgroups on two case studies of music-specific anhedonia, a condition in which listeners derive no pleasure from listening to music [28]. BW and NA are individuals who present with congenital and acquired music-specific anhedonia respectively. Both participants underwent a streamlined version of our study paradigm, with melodies presented 0, 4, 10, and 14 times, and only one melody per condition. We calculated the mean squared error (MSE) for liking and familiarity ratings of both the original and altered versions of these melodies using model predictions from the three-way (exposure X prediction error X music reward sensitivity) interaction models at all three levels of music reward sensitivity. As results of the mini meta-analysis indicated that there was no difference in the relationship between exposure and familiarity ratings across the music reward tertiles (i.e. no exposure X reward sensitivity interaction), we did not expect the model’s anhedonic predictions to have the lowest MSE for the familiarity ratings of our case studies. In contrast, since we did detect differences in the exposure-liking trajectory for musical anhedonics in our mini meta-analysis, we did expect the model’s predictions for anhedonics’ liking ratings to have the lowest MSE for our case studies.

For familiarity ratings, the model prediction at the hyperhedonic level best matched music-specific anhedonics’ responses (i.e. the model showed the lowest MSE of 18.13 at the hyperhedonic level), followed by the hedonic (19.95) and anhedonic (20.18) levels. This relatively better fit of the hyperhedonic predictions stems from the fact that all but one familiarity rating of the melodies that these participants made were extreme values of 1 or 6, thus likely representing a binary between knowing or not knowing these stimuli. This resulted in the lowest MSE for the hyperhedonic predictions, as the latter had the steepest (though not statistically significantly different) slope relating familiarity to exposure compared to the other tertiles. This suggests that both musical anhedonic cases were indistinguishable from hyperhedonics in their familiarity ratings, consistent with the finding that there were no significant interactions with music reward sensitivity from the meta-analysis of Studies 1-4 above. In contrast, for liking ratings, the model had the lowest MSE (3.66) from the anhedonic level when predicting the music-specific anhedonics’ data, compared to both the hedonic (4.97) and hyperhedonic (5.01) levels. This shows that the liking ratings of these cases were indeed more similar to the anhedonic group and different from that of the hedonics and hyperhedonics, consistent with the mini-meta analysis of Studies 1-4 above. Taking the familiarity and liking ratings together, these case studies provide further support for the idea that both cases of congenital and acquired musical anhedonia had less difficulty with learning these melodies than with deriving reward from them.

### Study 6

While the studies above are generally in support of the PC model, both schematic and veridical expectations were manipulated simultaneously, as the structural alterations introduced in the altered melodies violated both schematic and veridical expectations: schematic expectations since they contained statistically infrequent pitch patterns, but also veridical expectations as they violated participants’ specific predictions about that melody. Thus, to further probe whether the effect of prediction error in these studies can be attributed more to schematic or veridical expectation violation, we ran an additional follow-up study in which the melodies previously presented only in the post-exposure rating phase were now presented in the exposure phase (at 0, 2, 4, 8, and 16 times) and those originally in the exposure phase were now only presented in the post-exposure rating phase. Because the endings of the melodies in the exposure phase of this study are non-grammatical (meaning that there were less schematic expectations to be acquired for these endings), the prediction error manipulation is relatively limited to violations of veridical expectations. As a result, if subjective ratings are more sensitive to schematic expectations, then there should be no effect of prediction error in this study because the prediction errors do not violate a schematic expectation. Conversely, if these ratings are more sensitive to veridical expectations, then there should be an effect of prediction error, such that those that do not elicit a prediction error (i.e. presented in the exposure phase) are more familiar/preferred compared to those that do.

For familiarity ratings, we replicated the significant effect of exposure (β = 0.31, *t*(161) = 15.19, *p* < 0.001), but found no main effect of prediction error (β = 0.01, *t*(2129) = 0.48, *p* = 0.63) and no interaction between exposure and prediction error (β = -0.04, *t*(2786) = -1.77, *p* = 0.08) (Figure 3A). For liking ratings, we found a significant effect of exposure (β = 0.03, *t*(159) = 2.36, *p* = 0.02) and prediction error (β = 0.05, *t*(2595) = 2.27, *p* = 0.02), such that melodies that did not elicit prediction errors were preferred over those that did (Figure 3B). We did not detect an interaction between exposure and prediction error (β = -0.007, *t*(2519) = -0.31, *p* = 0.75). These results provide preliminary evidence that, while familiarity ratings were more sensitive to schematic expectations, liking ratings were more influenced by veridical expectations.

**Figure 3.**
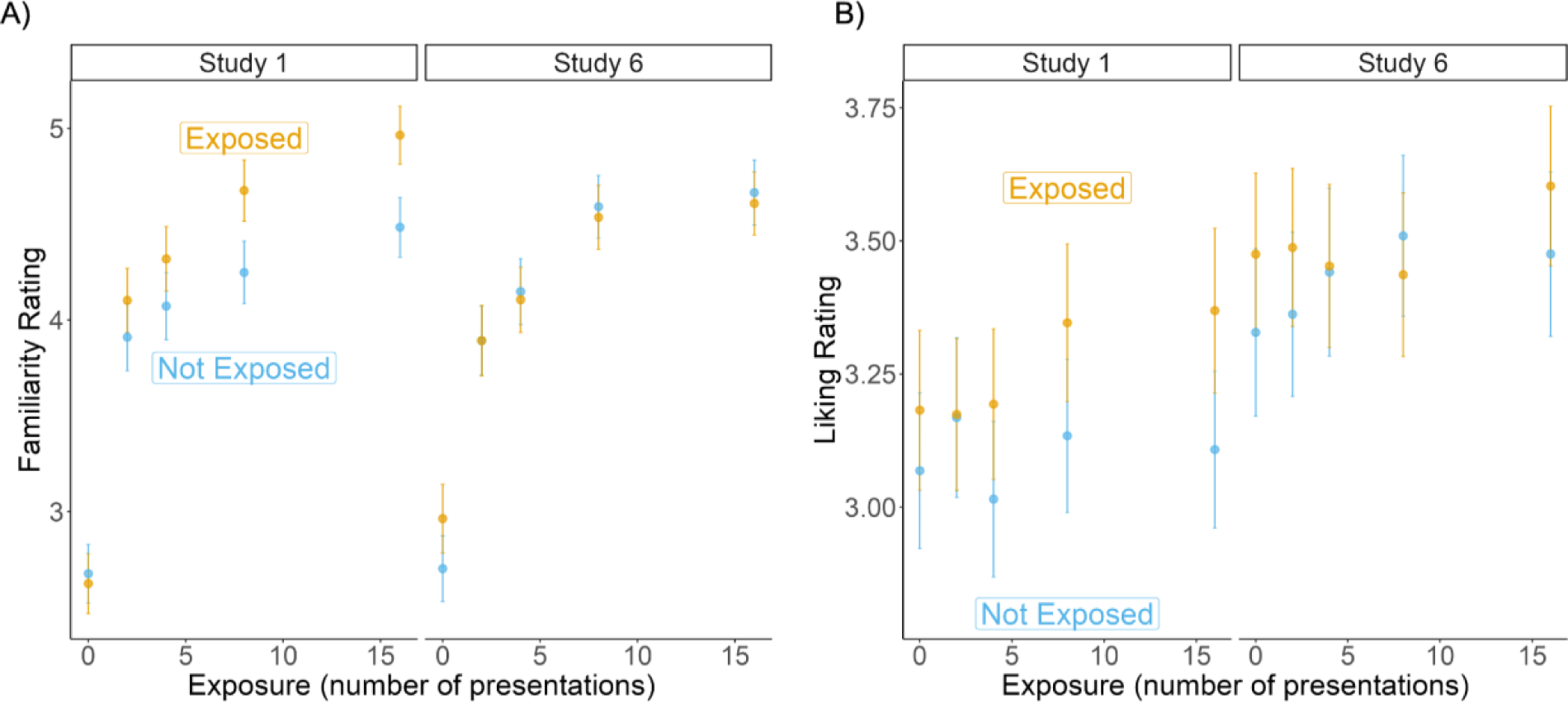
Mean and 95% confidence intervals of familiarity (A) and liking (B) ratings across Study 1 (exposed to fully grammatical melodies) and Study 6 (exposed to melodies with ungrammatical endings). “Exposed” melodies did not elicit prediction errors while “Not Exposed” melodies did.

### Mini Meta-Analyses of Studies 1 vs. 6

To further characterize the relationship between schematic and veridical expectations on familiarity and liking ratings, we aggregated data across Studies 1 (exposed to fully grammatical melodies) and 6 (exposed to melodies with ungrammatical endings). We then modeled both familiarity and liking ratings as a function of a three-way interaction between Study (1 vs. 6), prediction error, and exposure. This enables a direct comparison across studies in which there is a relative difference in the degree to which prediction errors violate schematic expectations (with there being more of a schematic expectation violation in the manipulation in Study 1). Thus, if ratings are more sensitive to schematic expectations, then there should be a significant study X prediction error interaction, such that the effect of prediction error is stronger for Study 1 compared to Study 6. Conversely, if ratings are more sensitive to veridical expectations, then the effect of alteration should be no different between these two studies.

#### Schematic expectations inform familiarity

For familiarity ratings, there was a significant three-way Study X prediction error X exposure interaction, such that the interaction between prediction error and number of presentations differed across the two studies (β = -0.14, *t*(5915) = -4.07, *p* < 0.001). While the effect of prediction error increased as a function of exposure in Study 1 (β = 0.1, *t*(5914) = 3.99, *p* = 0.001), the effect of prediction error remained the same across exposure in Study 6 (β = -0.04, t(5915) = -1.79, *p* = 0.07). Importantly, there was also a significant two-way Study X prediction error interaction, such that the effect of prediction error was weaker in Study 6 than in Study 1 (β = -0.15, t(5913) = -4.13, p < 0.001, mean and 95% confidence intervals shown in Figure 3A). There was no main effect of Study (β = 0.003, t(330) = 0.04, p = 0.97) nor a Study X exposure interaction (β = -0.02, t(327) = -0.96, p = 0.34). These results provide further evidence that familiarity is more sensitive to schematic expectations, and suggest that the relative increase in schematic expectations evoked by the melodies that did not elicit a prediction error in Study 1 was critical in increasing the effect of prediction error as a function of exposure.

#### Veridical expectations inform liking

For liking ratings, there was no significant three-way interaction between exposure, prediction error, and Study (β = -0.06, *t*(4192) = -1.82, *p* = 0.07) nor a two-way interaction between exposure and Study (β = 0.002, *t*(329) = 0.1, *p* = 0.92) or between prediction error and Study (β = -0.06, *t*(5163) = -1.81, *p* = 0.07). There was a main effect of Study on liking ratings, such that melodies were rated, overall, as more liked in Study 6 compared to Study 1 (β = 0.2, *t*(330) = 2.48, *p* = 0.01; mean and 95% confidence intervals shown in Figure 3B). As we did not detect a difference in the effect of prediction error across studies, these results suggest that, unlike familiarity ratings, liking ratings seem to be more informed by veridical expectations.

### Study 7

In the first 6 studies, participants were always asked to provide familiarity ratings before liking ratings. Because of this, one possible interpretation is that participants consistently rated the most familiar melodies from Studies 1-6 as most liked due to anchoring and/or demand effects. To rule out these possibilities, we ran an additional study in which participants completed the identical procedure as Studies 1 and 3, but did not rate any melodies on familiarity. There was still an effect of both exposure (β = 0.04, *t*(181) = 2.47, *p* = 0.01) and prediction error (β = 0.1, *t*(2227) = 4.87, *p* < 0.001) on liking ratings, but no interaction between the two (β = 0.001, *t*(2461) = 0.06, *p* = 0.95).

To formally compare whether removing familiarity ratings impacted the effect of our manipulations on liking ratings, we collapsed data from Studies 1 and 7 and modeled liking ratings as a three-way interaction between Study (1 vs. 7), prediction error, and exposure. This model did not detect any two-way interactions between Study and exposure (β = 0.008, *t*(349) = 0.36, *p* = 0.72) or Study and prediction error (β = -0.006, *t*(4310) = -0.21, *p* = 0.84), nor a three-way interaction (β = -0.05, *t*(3785) = -1.6, *p* = 0.11). Together, these results suggest the results of Studies 1-6 are not due to anchoring or demand effects.

### Study 8

Studies 1-7 together establish that effects of exposure and prediction error on liking and familiarity are not explained by task demands, and are blunted in groups with reduced reward sensitivity. Although the B-P scale is not used widely in any known culture, it is still possible that differences in the styles of music that we are exposed to from birth via our culture would impact how we learn and respond to these B-P melodies. To assess this possibility, we tested if the trajectories identified in Studies 1-7 are indeed similar across cultures. Study 8 extends the findings to investigate possible cultural effects on the process of becoming familiar with and preferring new pieces of music. To this end, we recruited 156 participants from China to complete the identical procedure as Study 4. For familiarity ratings, there was a significant interaction between exposure and prediction error (β = 0.08, *t*(1758) = 3.13, *p* = 0.002): the effect of prediction error (main effect β = 0.11, *t*(2437) = 4.49, *p* < 0.001) increased as a function of exposure (main effect: β = 0.3, *t*(154) = 14.9, *p* < 0.001). For liking ratings, we replicated both the significant main effect of exposure (β = 0.06, *t*(155) = 4, *p* = 0.001) and prediction error (β = 0.12, *t*(189) = 5.29, *p* < 0.001). There was no interaction between prediction error and exposure (β = 0.007, *t*(971) = 0.32, *p* = 0.75).

To further test whether familiarity and liking rating trajectories matched that of the US sample, we again fit two classes of models (logarithmic and linear for familiarity ratings; linear and quadratic for liking ratings) to these data. This revealed that, again, a logarithmic model best fit familiarity ratings (linear model AIC = 8958.3; logarithmic model AIC = 8376.9). A likelihood ratio test also indicated that a quadratic model fit the liking rating data better than a linear model (χ^2^(13) = 127.03, *p* < 0.001; see Figure 4 for model predictions plotted with mean and 95% confidence intervals), similar to the aggregated US sample.

**Figure 4.**
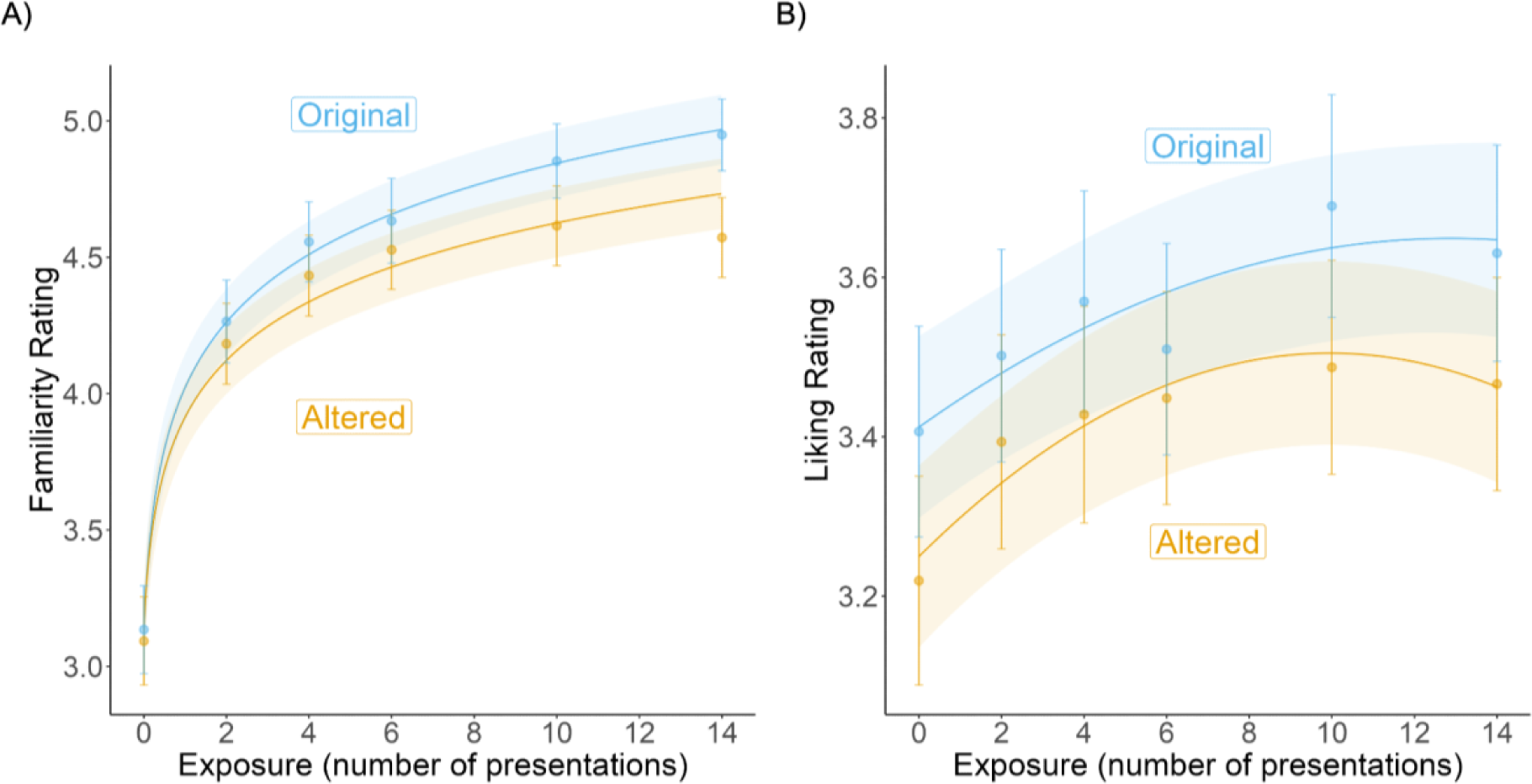
Cross-cultural replication of the effects of alterations and number of presentations on familiarity and liking ratings. Best fitting model predictions with mean ratings and 95% confidence intervals.

### Study 9

While the behavioral studies above provide support for cross-cultural applicability of the PC model, a key component of this model posits the involvement of the reward network in the brain. In Study 9, we relate exposure and prediction error to fMRI activity in the reward system. 21 young adults participated in the same study design as in Study 7 outside of the scanner, and then listened to the 8 melodies from Study 8 during fMRI as part of a larger-scale study in the lab looking at effects of music-based interventions in young adults and older adults [29]. Whole-brain, univariate analyses showed greater activation for melodies that did not elicit prediction errors compared to those that did in the right Heschl’s gyrus (Figure 5A), suggesting that the auditory cortex is sensitive to prediction errors.

**Figure 5.**
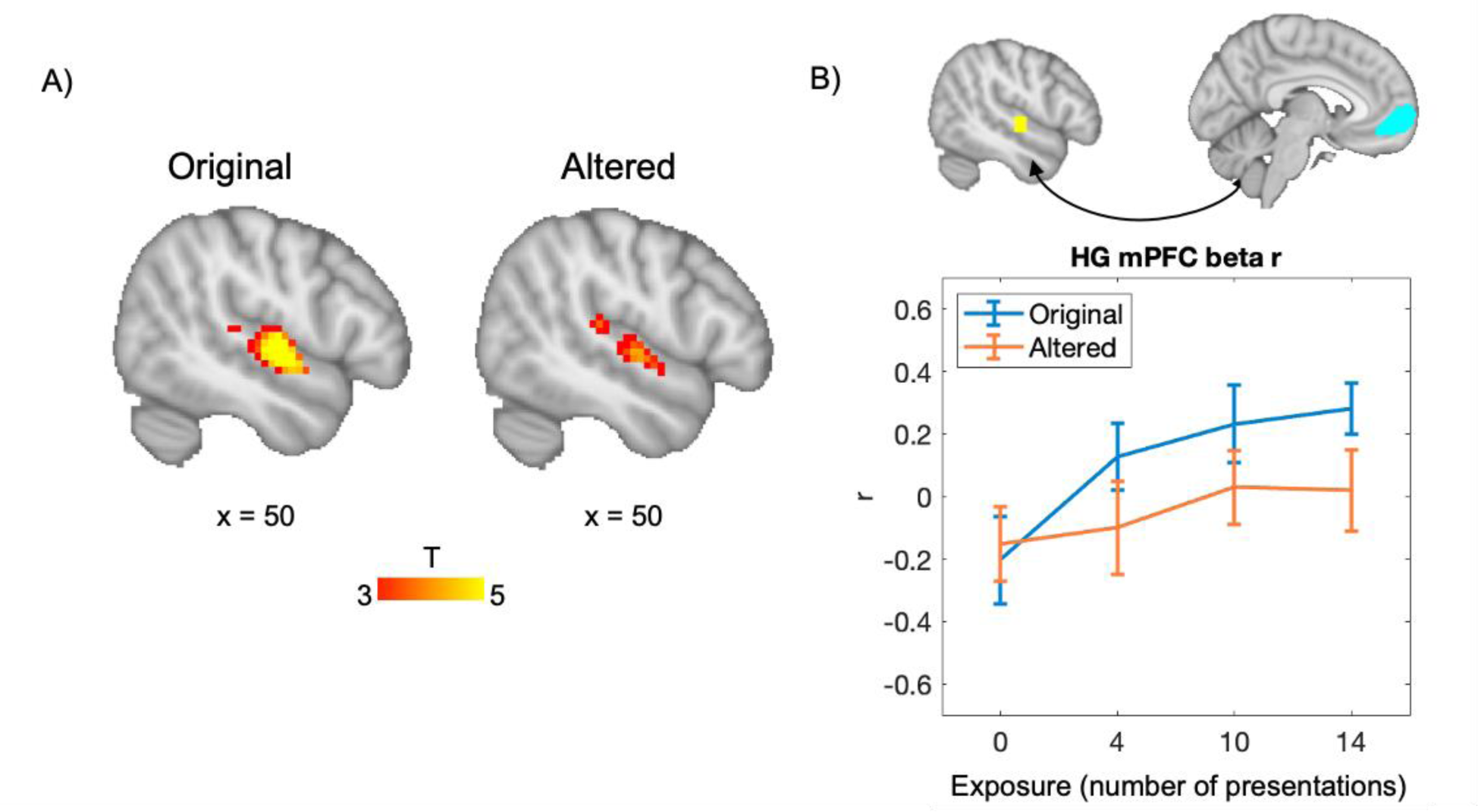
fMRI results. A) Greater activation for original (no prediction error elicited) than for altered (prediction error elicited) melodies in Heschl’s gyrus, confirming that auditory regions implement predictions. B) Higher functional connectivity, as quantified by correlations in beta-series, between auditory regions (Heschl’s gyrus) and reward regions (mPFC) for original melodies than for altered melodies (B) which increases with exposure (as quantified by number of presentations) for original but not for altered melodies.

Given previous evidence showing that co-activation of the reward and auditory brain areas is associated with musical preferences and learning [29], we also assessed the degree to which functional connectivity between these regions is modulated by predictive coding. The functional connectivity between auditory and reward areas was quantified by correlating the time series of beta-values extracted from Heschl’s gyrus (a sphere around the central voxel from the whole-brain analysis from Figure 5A) and reward-sensitive regions of interest (ROIs) in the nucleus accumbens and the medial prefrontal cortex (see Materials and Methods). A two-way within-subjects ANOVA with the dependent variable of auditory-reward functional connectivity, with the factors of prediction error and exposure, showed a significant main effect of prediction error (*F*(1,20) = 5.24, *p* = .033, ηp² = .21) and a significant main effect of exposure (*F*(3,60) = 3.31, *p* = .026, ηp² = .14). Figure 5B shows a linear relationship for original melodies as well as the effect of alteration. The same pattern was not observed for functional connectivity between Heschl’s gyrus and the nucleus accumbens (prediction error: *F*(1,20) = 1.61, *p* = .22, ηp² = .074; exposure: *F*(3,60) = .30, *p* = .83, ηp² = .015.)

## Discussion

Across nine studies, we provide novel evidence that clarifies the effects of both predictive coding and mere exposure on musical preference, by showing that listeners from two different cultures can rapidly learn from exposure and from prediction errors in novel music. This learning maps onto the brain’s reward system, and is sensitive to individual differences in reward sensitivity to music. In Studies 1-4, we established that changing the number of presentations (exposure) and altering the endings of melodies (prediction errors) both affected self-reported liking ratings for music, which ultimately provided evidence in support of predictive coding over mere exposure. Meta-analysis across Studies 1-4 and neuropsychological results from Study 5 confirmed that individuals with musical anhedonia formed familiarity in the same way as controls, but did not derive preferences from familiar sequences in the same way as their musically hedonic counterparts. Study 6 showed that familiarity ratings were more sensitive to schematic compared to veridical expectation violations, whereas liking ratings were more sensitive to veridical expectations. Study 7 tested for liking without testing for familiarity and established the same pattern of results, suggesting that effects of exposure and prediction error on liking were not due to anchoring effects. Study 8 established that both Chinese and American participants were affected by exposure and prediction errors in this musical system that was unfamiliar to both cultures. Finally, Study 9 ties this relationship between exposure, prediction error, and reward to increasing functional connectivity between the auditory and reward systems. Rather than simply showing that familiarity leads to liking, results reconcile the exposure-based and prediction-based accounts for music preference, and extend the PC model in three key directions: 1) towards unfamiliar, statistically and probabilistically novel music; 2) towards a more culturally-independent context via a cross-cultural comparison, and 3) towards its specific disruption in cases of musical anhedonia.

The degree to which veridical vs. schematic expectations influence musical reward has been difficult to assess, given that we are usually overexposed to particular musical genres that follow the same statistical patterns. The usage of the Bohlen-Pierce scale not only allows us to evade the pre-existing expectations that have accumulated from music exposure throughout our lives, but also offers greater insight into the relationship between veridical and schematic expectations. In Study 6, when the altered melodies were presented in the exposure phase, listeners preferred them now over their complements (which had been presented in the exposure phases of Studies 1 to 5). This indicates that the effect of prediction error in Studies 1-4 was not due to specific features of these melodies leading them to be more preferred.

Furthermore, participants’ ratings of liking but not familiarity continued to be responsive to the elicited prediction error in Study 6, suggesting a relative greater importance of veridical vs schematic expectations for musical preferences as there are less schematic expectations to use for learning the melodies in Study 6 compared to Study 1. Thus, it is possible that the greater veridical expectation learning in Study 6 (as compared to Study 1) additionally explained the main effect of study in our mini-meta analysis of Studies 1 and 6 on liking ratings. This mini meta-analysis also revealed a significant three-way interaction (study X prediction error X exposure) on familiarity ratings, such that the effect of prediction error increased as a function of exposure in Study 1 but not Study 6. We argue that this effect is due to the fact that the melodies presented during exposure in Study 6 evoked a relative decrease in schematic expectations while keeping veridical expectations the same relative to Study 1. Rather than claiming that familiarity always leads to liking, or that we only like what is familiar, the fact that schematic and veridical expectations differentially contributed to familiarity and liking ratings suggests that multiple, independent levels of prediction come into play in forming musical reward. The result is in line with prior work showing that repeated listening to a small number of B-P melodies (which increased veridical expectations without schematic expectations) resulted in higher preference ratings for those melodies, but non-repeated listening to a larger number of B-P melodies, while resulting in grammar learning (which is more akin to the learning of schematic expectations in the present study), did not lead to preference change [17].

Furthermore, both American and Chinese participants showed effects of both types of manipulations on preference ratings, clarifying the role of cultural background on the PCM model by suggesting that its influence is likely limited to situations in which there are differences in baseline implicit knowledge regarding the musical stimuli. On the other hand, individual differences on reward sensitivity appeared to play a crucial role in the process of linking predictive coding with musical reward. The consistency of the familiarity rating results (as well as the effect of prediction error on liking ratings) across tertiles underscores that musical anhedonics still learned the melodies, and were forming preferences to some degree. However, the fact that there was a difference in the effect of exposure on liking ratings across tertiles suggests that aesthetic preferences vary by the degree of exposure required to reach maximal preference. Specifically, because liking showed the strongest quadratic relationship with exposure in the anhedonic group, they may show less of an overall preference for music because they require less exposure before they become overexposed for their own liking.

Importantly, our study is the first to show that exposure to music de novo is associated with changes in the reward circuitry of the brain. Electrocortical (EEG and ECoG) recordings have shown that the middle Heschl’s gyrus is sensitive to melodic expectations [30], and fMRI studies have found that auditory and reward-related areas of the brain (including the amygdala, hippocampus, and ventral striatum) show increased activation during musical prediction errors [9] as well as during unexpected and/or unpredictable chord sequences [31]. However, as previous studies used familiar musical stimuli rooted in the Western musical tradition, it was not possible to determine when in the process of exposure the auditory and reward systems become engaged. Here, we observed that sensitivity to prediction errors emerge specifically in middle Heschl’s gyrus,thus extending previous EEG/ECoG results. Furthermore, increased functional connectivity between Heschl’s gyrus and mPFC was observed when listening to pieces that were more exposed, suggesting that the influence of repeated exposure on liking is subserved by changes in communication between the auditory and reward networks.

Several outstanding questions stem from these studies that warrant future exploration. First, it remains to be seen whether preference ratings would continue to increase with more than 16 exposures. It is quite possible that the positive relationships found here between exposure and liking is reflective of the positive side of a quadratic function, and that if we were to extend the number of repetitions in this paradigm, we would see preference ratings begin to decrease at an inflection point. Given that we chose to optimize for longer, more dynamic pieces of music, it was not feasible to increase the number of exposures beyond 16 without altering other key aspects of the design, introducing fatigue or habituation, or otherwise increasing cognitive demand in ways that would confound the study. Future studies with shorter stimuli may be able to assess the full extent of the relationship between liking and repetition in B-P stimuli and the degree to which relative frequencies (14 relative to 10 vs 14 relative to 2) play a part.

Second, while the current fMRI study shows sensitivity to prediction in the reward system, it is not sufficiently powered to assess possible individual differences in neurobiology between musical anhedonics and hedonics. Previous neuroimaging studies that included participants with musical anhedonia have shown reduced structural and functional connectivity between auditory and reward-sensitive areas in musical anhedonics [32, 33], and that alterations of fronto-striatal pathways can lead to either increases or decreases in subjective liking ratings of music [34]. Future neuroimaging studies are needed in this special population, and also across cultures, to establish how the mechanisms of learning relate to auditory-reward connectivity. In sum, we developed an innovative paradigm to assess the effects of exposure and prediction errors in novel music on musical preference across cultures and in special populations. Our results are the first to show the multiple levels by which exposure and prediction errors in music generate reward, and provide strong evidence for this learning process across two cultures.

Individuals with musical anhedonia did not show the same patterns as a result of exposure, offering a testable mechanism by which the human brain learns to predict sounds from our environment and to map those predictions onto reward. As the relationship between predictions and reward underlie much of motivated behavior [7, 8, 18], examining the emergence of this relationship during the course of a study may provide a better understanding of how these foundational neurocognitive systems may go awry in a variety of psychiatric and neurological disorders.

## Materials and Methods

### Stimuli

The stimuli used in all studies were composed in the Bohlen-Pierce Scale. While most musical systems around the world are based around the octave, which is a 2:1 ratio in frequency, the B-P scale is based on a 3:1 ratio (*tritave* rather than octave) that is divided into 13 logarithmically even steps. This 13-tone scale can be used to generate musical intervals and chords which have low-integer ratios and are perceived as psychoacoustically consonant (Mathews, 1988). While music in B-P scale is known to some composers, performers, conductors and scholars, it is considered “non-standard music” (35) and has not been adopted into any mainstream musical culture to date. Monophonic melodies were composed in the B-P scale by a musician and research assistant in the lab (E.Z.) in the digital audio workstation Ableton Live on a Korg nanoPAD2 USB MIDI and played on a MIDI clarinet instrument from the plugin library Xpand!2 by Air Music Tech. The clarinet was chosen because its timbre has higher energy at odd harmonics than at even harmonics; this spectral distribution is easier to learn due to its congruence with the B-P scale [22]. In total, 14 20s Bohlen-Pierce melodies were composed that followed the same artificially-derived harmonic structure from past studies [17]. Light compression and reverb were applied to all stimuli to bring them to the same volume, and were subsequently exported as 44.1kHz .mp3 files. To generate melodies that contained an error in local prediction, an altered version of each melody was also created, which was identical to the original piece except for the ending, which was changed to violate the musical structure of the B-P scale. Specifically, violations consisted of deviations from the chordal tones of the last chord [17, 36, 37], such that they disrupt the harmonic structure of the established melody. The original and altered melodies are available online at https://osf.io/n84d5/, along with the preregistration as well as data and code for this study. In all studies except Study 6, the altered melodies were presented only once, during the post-exposure phase. Finally, two of the melodies were used only as part of the perceptual cover task (during the exposure phase). A vibrato effect was added to a single note in these two melodies and during the task, participants were asked to press a key whenever they heard the vibrato note. To decrease expectations, we created six versions of each, where the location of this vibrato note varied across each version.

### Study 1

#### Participants

A priori power analysis using pilot data (n = 46) indicated that a sample size of 165 would achieve 0.80 power to detect a medium effect size (Cohen’s *f* = 0.27) of the effect of the number of presentations on liking ratings at a significance level of 0.05. Participants were Prolific workers in the United States between the ages of 18-65. We recruited 234 participants for Study 1, of which 66 participants were excluded for failing our perceptual cover task (see Procedure below), resulting in a final sample size of N = 169 (104 female; mean age = 32.03).

To measure individual differences in music reward sensitivity and identify musical anhedonics, participants completed the BMRQ, a 20-item questionnaire based on five factors: musical seeking, emotion evocation, mood regulation, sensory-motor, and social reward. Participants also completed the Goldsmith Musical Sophistication Index (Gold-MSI), a self-report measure of musical skills and behaviors [38], the Revised Physical Anhedonia Scale (PAS), a self-report measure of general anhedonia [39], and the Ten-Item Personality Inventory (TIPI), a brief measure of the Big-Five personality traits [40]. All scales were scored in accordance with the original publications.

#### Procedure

After consenting, participants were screened using an online headphone check [41] to ensure that they were using headphones and could hear our stimuli properly before undergoing the three phases of our study. In phase 1 (pre-exposure), participants listened to 8 of the B-P melodies, one at a time, and provided liking ratings, using a Likert-scale ranging from 1 (‘strongly dislike’) to 6 (‘strongly like’) and familiarity ratings, from 1 (‘not familiar at all’) to 6 (‘very familiar’) for each melody. As the pre-exposure ratings are intended for a different analysis on the effects of novelty rather than reward learning, they will be presented in a separate report; here we focus on post-exposure ratings.

In phase 2 (exposure), the 8 melodies heard in phase 1 were played for participants a varying number of times (either 2, 4, 8 or 16 with two melodies in each condition). The specific melodies in each of the 4 exposure conditions was counterbalanced across participants. Furthermore, the presentation order was pseudorandomized so that no melody was heard consecutively. During this phase, participants were asked to complete a perceptual cover task, in which they were instructed to listen for notes that contained a “warble” sound (vibrato) and to press the “v” key on their keyboard as soon as they heard one. Six of the trials (created from two different B-P melodies) heard in the exposure phase contained vibrato notes, with the vibrato occurring at different points of the melody. In total, participants heard 66, 20s melodies during phase 2, resulting in an exposure phase that lasted 22 minutes.

During phase 3 (post-exposure), participants heard each of the 8 melodies again (without vibrato), along with 2 new melodies that they had not heard in phase 1 or 2 (0 presentation condition) as well as the altered versions (different endings) of these ten melodies. Participants provided liking and familiarity ratings for each of these 20 trials, using the same scale as in phase 1.

After completing phase 3, participants were redirected to an online survey where they provided demographic information and completed individual difference measures including the BMRQ and PAS.

#### Exclusion criteria

Participants who did not accurately perform the surface task of identifying the warble/vibrato notes during exposure were removed from all subsequent analyses. Specifically, for each participant, we calculated d-prime from the total number of hits (number of vibrato melodies for which a ‘v’ was pressed), misses (number of vibrato melodies for which a ‘v’ was not pressed), false alarms (number of vibrato melodies for which a ‘v’ was not pressed) and correct rejections (number of non-vibrato melodies for which a ‘v’ was not pressed). D-prime was calculated from the difference between z-transformed hit and false-alarm rates, with the adjustment where 0.5 errors were assumed for participants who made no errors [42]. The d-prime measure therefore indicates how well participants could discriminate between a warble note and a non-warble note and was used to remove participants who did not follow instructions for the surface task. Any participant who had a d-prime measure of less than 1 was removed from subsequent analyses [42], as was specified in our pre-registration. However, in follow-up analyses we did explore whether keeping the participants who did not reach the d-prime criterion changed the results; these exploratory analyses are included in Supplementary Materials.

### Study 2

#### Participants

To maintain consistency, we used the same target sample size from our a priori power analysis for Study 1 for Studies 2-4. We recruited 221 participants. 57 participants were excluded for failing a perceptual cover task, resulting in a total sample size of 164 (93 female, mean age = 32.67).

#### Procedure

Participants underwent the same procedure as in Study 1, with the exception that 10 melodies were presented either 0, 2, 4, 6, 10, or 14 times during the exposure phase (2 melodies in each condition).

### Study 3

#### Participants

We recruited 214 participants, 45 of whom were excluded for failing our perceptual cover task, resulting in a total sample size of 169 (89 female; mean age = 32.27).

#### Procedure

Participants underwent the exact same procedure as in Study 1, with the exception that the order of melodies heard in the pre-exposure phase was completely randomized.

### Study 4

#### Participants

We recruited 222 participants, 57 of whom were excluded for failing our perceptual cover task, resulting in a total sample size of 165 (83 female; mean age: 31.78).

#### Procedure

Participants underwent the exact same procedure as in Study 2, with the same 10 melodies during exposure phase, with the exception that the order of melodies heard in the pre-exposure phase were randomized and counterbalanced across participants.

### Study 5

#### Participants

The congenital music specific anhedonic (initials BW, 58-year-old male) had participated in a previous case study in our lab [33]. The acquired music specific anhedonic (initials NA, 53-year-old female) had reached out to the final author after self-reporting a loss in pleasure derived from music listening after having received rTMS treatment for depression after the death of a loved one. As both of these cases were self-identified as musically anhedonic, rather than recruited online using Prolific, they were treated as separate case studies rather than included in the same group for Studies 1 through 4. Both of these cases had low scores on the extended BMRQ (eBMRQ BW = 30; NA = 43; [43] but normal PAS scores (PAS-auditory: BW = 8, NA = 4; PAS-non-auditory: BW = 14, NA = 15).

#### Stimuli

We used a subset of four non-altered melodies which were rated, on average, the highest in post-exposure liking ratings across Studies 1-4 for Study 6. These, along with their altered versions, resulted in eight unique melodies presented to participants in this study. Participants also completed an updated version of the BMRQ: the extended Barcelona Music Reward Questionnaire (eBMRQ), which includes an additional sixth factor consisting of 4 additional items which measures experiences of absorption in music listening [43].

#### Procedure

Participants underwent the same procedure as previous studies, with the exception that melodies were presented either 0, 4, 10, or 14 times during the exposure phase and that there was only one melody assigned to each condition.

### Study 6

#### Participants

We recruited 279 participants, 116 of whom were excluded for failing our perceptual cover task, resulting in a total sample size of 163 (64 female; mean age: 35.46).

#### Procedure

Participants completed the same procedure as in Study 1, with the exception that altered melodies were presented in the pre-exposure and exposure phase of the study. In this study, original melodies were only presented in the post-exposure phase.

#### Stimuli

The same stimuli used in Studies 1 and 3 were used in Study 5. Participants in Study 5 also completed the eBMRQ instead of the BMRQ.

### Study 7

#### Participants

We recruited 244 participants, 64 of whom were excluded for failing our perceptual cover task, resulting in a total sample size of 180 (78 female; mean age = 35.62).

#### Procedure

Participants underwent the exact same procedure as in Study 1, with the exception that they were not asked to provide familiarity ratings.

### Study 8

#### Participants

Participants were recruited via WeChat, a Chinese instant messaging app. A poster containing a QR code was sent in several group messages of students of Beijing Normal University, who subsequently shared this code via word of mouth and personal WeChat messages. We recruited 216 participants. 56 were excluded for failing our perceptual cover task and 4 for completing the task twice, for a total of 156 (106 female; mean age: 23.09).

#### Stimuli

The same stimuli used in Studies 2 and 4 were used in Study 7. Participants in Study 7 also completed the eBMRQ instead of the BMRQ.

#### Procedure

The QR code led to a questionnaire that recorded participants’ name and email address. An email was then sent to the address participants provided, which contained a link to the experiment. This link redirected participants to our experiment, in which they subsequently underwent the same Procedure as Study 3.

### Study 9

#### Participants

Participants in this study were either undergraduates at Northeastern University who completed the study (both the online task and an in-person fMRI scan) for course credit or young adults recruited via word-of-mouth from the Boston area. A total of 21 participants (15 female, mean age = 19.8) completed the fMRI version of our task.

#### Stimuli

The same stimuli and materials that were used in Study 6 were used in Study 7, including the eBMRQ.

#### Procedure

Participants underwent the same procedure as in Study 6 as well as an fMRI scan immediately after completing the online behavioral study. During the scan, participants listened to 24 clips of music once. Eight of the clips were Bohlen-Pierce melodies that participants had heard previously during the task (at 0/4/10/14 presentations; both original and altered melodies). The remaining trials acquired were not in the Bohlen-Pierce scale and were not used in the analysis for the present study. Each trial consisted of 20s of passive listening, followed by 2s to rate the melody for liking (on a scale of 1-4), and 2s to rate the melody for familiarity (also 1-4 scale).

#### fMRI Data Acquisition

Images were acquired using a Siemens Magnetom 3T MR scanner with a 64-channel head coil at Northeastern University Biomedical Imaging Center. fMRI data were acquired as echo-planar imaging (EPI) functional volumes covering the whole brain in 48 axial slices (fast TR = 475 ms, TE = 30 ms, flip angle = 60°, FOV = 240mm, voxel size = 3 × 3 × 3 mm^3^, slice thickness = 3 mm, anterior to posterior, z volume = 14.4 mm) in a continuous acquisition protocol of 1440 volumes for a total acquisition time of 11.4 minutes. T1 images were also acquired using a MPRAGE sequence, with one T1 image acquired every 2400 ms, for approximately 7 minutes. Sagittal slices (0.8 mm thick, anterior to posterior) were acquired covering the whole brain (TR = 2400 ms, TE = 2.55 ms, flip angle = 8°, FOV= 256, voxel size = 0.8 × 0.8 × 0.8 mm^3^). As part of the existing protocol we also acquired resting state and DTI sequences, but these were not used for this study.

#### fMRI Data Analysis

*Pre-processing.* fMRI data were preprocessed using the Statistical Parametric Mapping 12 (SPM12) software [44] with the CONN Toolbox [46]. Preprocessing steps included functional realignment and unwarping, functional centering, slice time correction, outlier detection using the artifact detection tool, functional and structural segmentation and normalization to MNI template, and functional smoothing to an 8mm gaussian kernel [46]. Denoising steps for fMRI data included white matter and cerebrospinal fluid confound correction [47], and bandpass filtering to 0.008– 0.09 Hz.

*First-level analysis.* First- and second-level analyses were completed in SPM12. For each participant, data were converted from 4D to 3D images, resulting in 1440 scans. The model was specified using the following criteria: interscan interval = 0.475 seconds, microtime resolution = 16, microtime onset = 8, duration = 42. Only data from the time while the participant was listening to the musical excerpt were included in this model. Each of the 8 trial types (0/4/10/14 presentations of both original and altered melodies) was modeled separately, and trials during which participants were listening to non-BP melodies were included as a separate condition so as to be regressed out of the model’s intercept. The resulting first-level contrasts were then analyzed using a one-sample t-test across all participants at the second level. Whole-brain results were rendered to a standard MNI brain. Results from the second-level analyses were statistically corrected using a voxel threshold of p < 0.05 (FDR-corrected) through CONN Toolbox. For Beta-weights for ROIs in the Heschl’s gyrus (HG) and medial prefrontal cortex (mPFC) were extracted from participants’ first-level SPM.mat files using the CONN Toolbox atlas and correlated separately for each trial to test for the effects of alteration and number of presentations on the functional connectivity between auditory and reward-sensitive regions.

