## Supplemental materials for "Generating New Musical Preferences from Multi-level Mapping of Predictions to Reward"

for

**Generation of New Musical Preferences from Hierarchical Mapping of Predictions to Reward**

### Supplementary Results

#### *Reanalysis of Studies 1-4 with Excluded Participants*

Although the pre-registered analysis in the main manuscript excluded participants who did not reach a d-prime criterion on the cover task during phase 2, as an exploratory analysis we re-ran all of our analyses for Studies 1-4 separately including participants who were excluded. All of our results remained the same except for the detected number of presentations X alteration interaction on liking ratings in Study 1, which became insignificant ($\beta = 0.03$, $t(2423) = 1.38$, $p = 0.17$).

#### *Meta Analyses of Studies 1-4*

<u>Musical experience additionally influences exposure-liking trajectory</u>

To determine the potential impact of individual differences on music training and engagement on reward learning, we split our sample from Studies 1-4 based on responses on the Goldsmiths Musical Sophistication Index (GoldMSI general sophistication score: Low: 59-74; Average: 75-79; High: 80-99). We then added this variable (which was dummy-coded to treat the average group as the reference level) as an interaction term to our best fitting models (logarithmic for familiarity ratings; quadratic for liking ratings). We interpreted any interaction between number of presentations and general sophistication score from the GoldMSI as evidence that the relationship between familiarity and/or liking ratings and number of presentations differed across groups.

For familiarity ratings, we did not detect any differences in ratings across the three tertiles. Further, there were no significant two-way interactions between general sophistication and number of presentations or alteration. No three-way interaction between general sophistication, number of presentations, and alteration was significant (see Figure S1).

For liking ratings, we again did not detect any differences in ratings across the three tertiles. However, we did detect a significant two-way interaction between the linear effect of number of presentations and general sophistication score ($\beta = 0.07$, $t(649) = 2.56$, $p = 0.01$): the linear effect of number of presentations on liking ratings was stronger for those with high general sophistication scores compared to those with average scores relative to our sample. We did not detect any significant three-way (melody alteration X number of presentations X general sophistication score) interactions (see Figure S1).

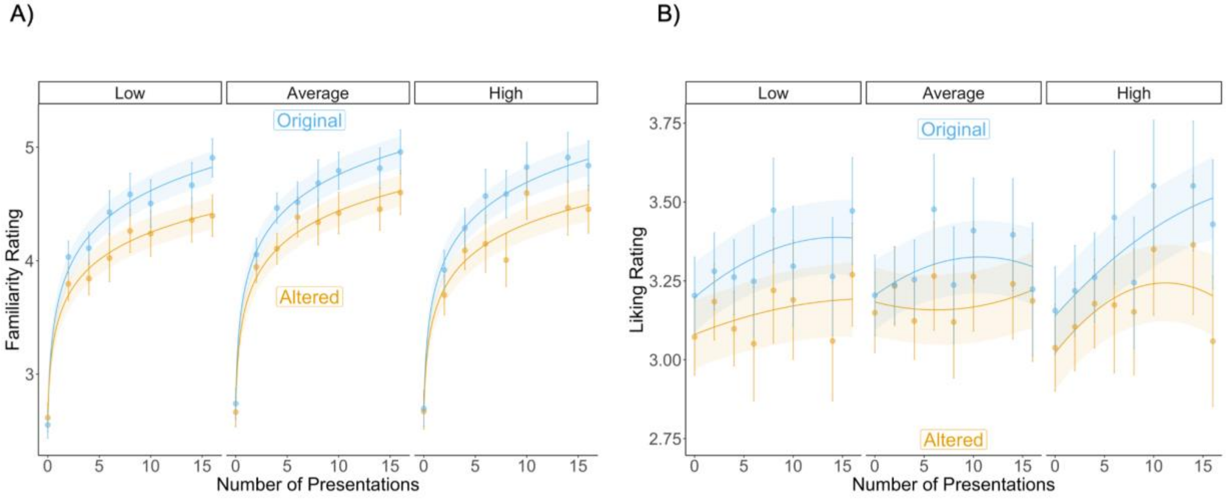


**Figure S1:** *Model-predicted familiarity (A) and liking (B) ratings as a function of number of presentations, alteration, and general musical sophistication score (tertile split on general sophistication score of the GoldMSI: low, average, and high groups). Points and associated error bars represent 95% confidence intervals.*
